# Reassessing global historical ℛ_0_ estimates of canine rabies

**DOI:** 10.1101/2024.04.11.589097

**Authors:** Michael Li, Michael Roswell, Katie Hampson, Benjamin M. Bolker, Jonathan Dushoff

## Abstract

Rabies spread by domestic dogs continues to cause tens of thousands of human deaths every year in low- and middle-income countries. Nevertheless rabies is often neglected, perhaps because it has already been eliminated from high-income countries through dog vaccination. Estimates of canine rabies’s intrinsic reproductive number (*ℛ*_0_), a metric of disease spread, from a wide range of times and locations are relatively low (values *<* 2), with narrow confidence intervals. Given rabies’s persistence, this consistently low and narrow range of estimates is surprising. We combined incidence data from historical outbreaks of canine rabies from around the world with in-depth contact-tracing data from Tanzania to investigate initial growth rates (*r*_0_), generation-interval distributions (*G*), and reproductive numbers (*ℛ*_0_). We improved on earlier estimates by choosing outbreak windows algorithmically; fitting *r*_0_ using a more appropriate statistical method that accounts for decreases through time; and incorporating uncertainty from both *r*_0_ and *G* in our confidence intervals on *ℛ*_0_. Our *ℛ*_0_ estimates are larger than previous estimates, with wider confidence intervals. These revised *ℛ*_0_ estimates suggest that a greater level of vaccination effort will be required to eliminate rabies than previously thought, but that the level of coverage required remains feasible. Our hybrid approach for estimating *ℛ*_0_ and its uncertainty is applicable to other disease systems where researchers estimate *ℛ*_0_ by combining data-based estimates of *r*_0_ and *G*.

## Introduction

Canine rabies, primarily spread by domestic dogs, is a vaccine-preventable disease that continues to cause tens of thousands of human deaths every year in low- and middle-income countries (LMICs) (Taylor et al., 2017; Minghui et al., 2018). Canine rabies has been eliminated from high-income countries by mass dog vaccination (Rupprecht et al., 2008). Despite the effectiveness of dog vaccination, rabies continues to cause many human deaths and large economic losses in LMICs due to the limited implementation of rabies control strategies (Hampson et al., 2015). The past two decades have seen an increase in rabies control efforts, including dog vaccination campaigns and improvements in surveillance (Kwoba et al., 2019; Mtema et al., 2016; Gibson et al., 2018; Mazeri et al., 2018; Wallace et al., 2015). The World Health Organization (WHO) and partners (OIE, FAO, GARC) joined forces to support LMICs in eliminating human deaths from dog-mediated rabies by 2030 (Minghui et al., 2018; Abela-Ridder et al., 2016). Mass dog vaccination campaigns have begun in some LMICs and are being scaled up (Castillo-Neyra et al., 2019; Evans et al., 2019). However, the emergence of SARS-CoV-2 pandemic disrupted rabies control and elimination efforts (Nadal et al., 2022). As the SARS-CoV-2 pandemic is transitioning out of global emergency, rabies control programmes are resuming. An understanding of rabies epidemiology — in particular, reliable estimates of the basic reproductive number (*ℛ*_0_), a quantitative measure of disease spread that is often used to guide vaccination strategies — could inform rabies control efforts.

The basic reproductive number *ℛ*_0_ is defined as the expected number of secondary cases generated from each primary case in a fully susceptible population (Macdonald, 1952). Estimates of *ℛ*_0_ for rabies have been made using various methods including direct estimates from infection histories, epidemic tree reconstruction, and epidemic curve methods. These *ℛ*_0_ estimates based on historical outbreaks of rabies that span a variety of regions and time periods have generally been surprisingly low, typically between 1 and 2 with narrow confidence intervals (Hampson et al., 2009; Kurosawa et al., 2017; Kitala et al., 2002). With such a low *ℛ*_0_ one might expect rabies to fade out due to a combination of behavioural control measures and stochastic fluctuations, even in the absence of vaccination. In contrast to diseases that have already been eradicated, but that have a large *ℛ*_0_ (e.g., rinderpest, with *ℛ*_0_ *≈* 4 (Mariner et al., 2005)), *ℛ*_0_ estimates for rabies suggest that control through vaccination should be relatively easy.

Here we revisit and explore why rabies, with its low *ℛ*_0_, nonetheless persists in many countries around the world. Such persistence suggests that rabies’s potential for spread, and therefore the difficulty of rabies control, may have been underestimated. In this paper, we combine information derived from epidemic curves with a highresolution contact tracing data set that provides large numbers of observed generation intervals (which is rare for infectious disease studies) to estimate *ℛ*_0_.

## Materials and Methods

*ℛ*_0_ is often estimated by combining two other epidemiological quantities: the initial growth rate of an epidemic (*r*_0_) and the generation interval (*G*) distribution, where the generation interval is defined as the time between successive infections along a transmission chain (Park et al., 2018). The initial growth rate *r*_0_ is often estimated by fitting a model to time series data from the early stages of epidemics. *G* is an individual-level quantity that measures the time between an individual getting infected to infecting another individual. The generation interval distribution is the natural way to link *r*_0_ and *ℛ*_0_ (Wallinga and Lipsitch, 2006; Champredon and Dushoff, 2015). *ℛ*_0_ can be estimated from *r*_0_ and the *G* distribution by the Euler-Lotka equation (Wallinga and Lipsitch, 2006)

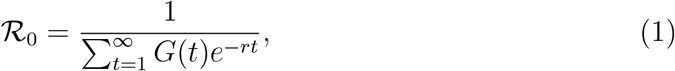

where *t* is time, and *G*(*t*) is the generation interval distribution. This formula is convenient to calculate point estimates of *ℛ*_0_; however, researchers rarely propagate uncertainty from the estimates of *r*_0_ and the *G* distribution through this formula.

### Initial growth rate

Disease incidence typically increases approximately exponentially during the early stages of an epidemic. The initial growth rate *r*_0_ is often estimated by fitting exponential curves from near the beginning to near the peak of an epidemic. However, growth rates estimated from an exponential model can be biased downward, overconfident, and sensitive to the choice of fitting windows (Ma et al., 2014). Here we used logistic rather than exponential curves to more robustly estimate *r*_0_ (Ma et al., 2014; Chowell, 2017).

We selected fitting windows algorithmically for each outbreak as follows: (1) we break each time series into “phases”: a new phase starts after a peak with a height of at least `minPeak` (16) for this MS) cases, followed by a proportional decline in cases of at least `declineRatio` (0.25); (2) In each phase, we identify a prospective fitting window starting after the last observation of 0 cases and extending one observation past the highest value in the phase (unless the highest value is itself the last observation); (3) we then fit our model to the cases in the fitting window if (and only if) it has a peak of at least `minPeak` cases, a length of at least `minLength` (4) observations, and a ratio of at least `minClimb` (1.5) between the highest and lowest observations. We tried a handful of parameter combinations before settling on a final set during an expert consultation. These explorations are detailed in our code repository.

### Observed Generation intervals

An earlier rabies study constructed generation intervals by summing two quantities: a latent period (the time from infection to infectiousness), and a wait time (time from infectiousness to transmission) (Hampson et al., 2009). Since clinical signs and infectiousness appear at nearly the same time in rabies, the incubation period (the time from infection to clinical signs) is routinely used as a proxy for the latent period. Hampson et al. randomly and independently resampled latent (really, incubation) periods and infectious periods from empirically observed distributions (Hampson et al., 2009), and then sampled waiting times uniformly from the selected infection periods. However, constructing *G* values by summing independently resampled values of incubation and infectious periods accounts neither for the possibility of multiple transmissions from the same individual, nor for correlations between time distributions and biting behaviour. Figure 1 illustrates the generation intervals of a single transmission event from a rabid animal (comprising a single incubation period plus a waiting time) and multiple transmission events from a rabid animal (comprising a single incubation period and three waiting times). In cases where transmission links are not directly observed, one should consider reweighting incubation-period observations to account for unequal transmission from different infectors. In our case, we can account for these effects directly by relying on generation intervals observed through contact tracing.

**Figure 1:**
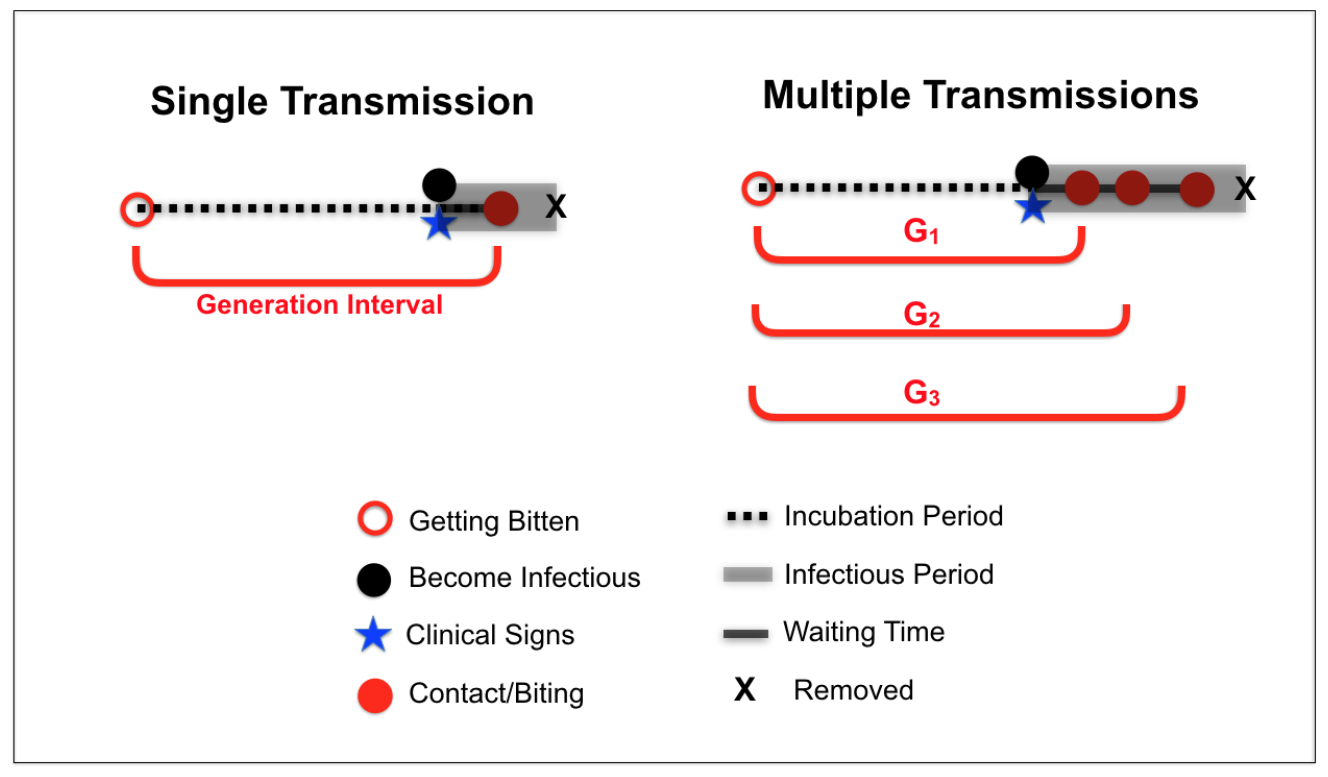
Decomposing generation intervals. Generation intervals start when a focal animal acquires infection (open red circle) and end after a period of viral replication (dashed line) when an animal shows clinical signs (blue star), becomes infectious (solid black circle) and infects another animal — in rabies, the onset of clinical signs and of infectiousness are closely synchronized. Once the infectious period (grey block) starts, there is a wait time (solid black line) until a susceptible host (solid red circles) is bitten. The infectious period ends with the death of the focal host (black X). The generation interval is the interval between the focal animal getting infected, and when it infects a new case (red interval between open and solid circles). If a single biter transmits multiple times (right), the wait times generally vary, but the incubation period is the same for each transmission event.

### Correcting for vaccination

In a population where some animals are not susceptible, calculations based on estimates of *r*_0_ and the *G* distribution (1) do not estimate *ℛ*_0_, but instead estimate the *realized* average number of cases per case, also known as the effective reproductive number *ℛ*_e_. In the case of rabies, vaccination is the only known cause of immunity (case fatality in dogs is believed to be 100%). For a given population with *?* vaccination proportion, the estimated *ℛ*_0_ with correction for vaccination is

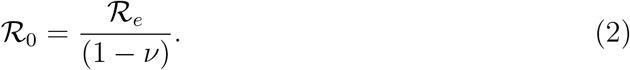

### Data

We used data from December 2002 – November 2022, from an ongoing contact tracing study in Tanzania (Hampson et al., 2008, 2009). The data set contains 8636 domestic dog recorded events (i.e., domestic dogs bitten by an animal), and 3552 suspected rabid dogs in the Serengeti District of northern Tanzania. Transmission events were documented through retrospective interviews with witnesses, applying diagnostic epidemiological and clinical criteria from the six-step method (Tepsumethanon et al., 2005). Each dog was given a unique identifier, and date of the bite and clinical signs were recorded if applicable and available. 2132 of the dog transmissions were from unidentified domestic animals or wildlife. We restricted our analysis to domestic dog transmissions (i.e., dog to dog), and obtained 293 directly observed generation intervals (i.e. both biter and secondary case have “time bitten” records). There were four observed dogs with multiple exposures (i.e., bitten by different identified biters), generating extra generation intervals, but it is unclear which transmission event transmitted rabies to these dogs. For simplicity, we omitted these four dogs and their generation intervals from our analysis.

### Fitting and Propagating Parameter Uncertainties

To propagate uncertainties for both *r*_0_ and *G*, we used a hybrid approach. We first fit logistic models, with negative binomial observation error, to incidence data to estimate *r*_0_ implemented in the R package `epigrowthfit` Jagan et al. (2024). We then compute a sample of 1000 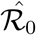 values using equation (2). For each value of 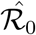, we first draw a value of 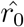 from a Normal distribution matching the estimated sampling distribution of the logistic fit parameters and an independent sample of *G* from the empirical contact tracing data. To sample *G* from the empirical contact tracing data, we first take a weighted sample of 100 biters, which accounts for biter-level variation, and for each biter, we sample a *G* from its respective transmission event, to account for individual variation. We then matched samples of *G* to the *r*_0_ samples to produce a range of estimates for *ℛ*_0_. This hybrid sampling approach incorporates the uncertainties in both *r*_0_ and *G* into the distribution of *ℛ*_0_ estimates. Finally, we use the 2.5, 50, and 97.5 percentiles of the distribution of *ℛ*_0_ estimates to get point estimates and confidence limits for *ℛ*_0_ for each rabies outbreak.

## Results

Figure 2 shows the empirical distributions of the observed incubation periods, rabid dog biting frequency, and generation intervals from the contact tracing data. The mean observed incubation period is 27.5 days (*n* = 1109 dogs), the mean biting frequency is 1.65 bites per rabid dog, and the weighted mean incubation period is 36.6 days (*n* = 143 biting dogs). The mean observed generation interval is 37.9 days (*n* = 143 primary infections resulting in 293 secondary cases), which is substantially larger than the mean generation interval constructed from summing independently sampled incubation periods and wait times (24.9 days (Hampson et al., 2009)). The weighted incubation period distribution is a better approximation of the generation interval distribution than the unweighted incubation period of all dogs.

**Figure 2:**
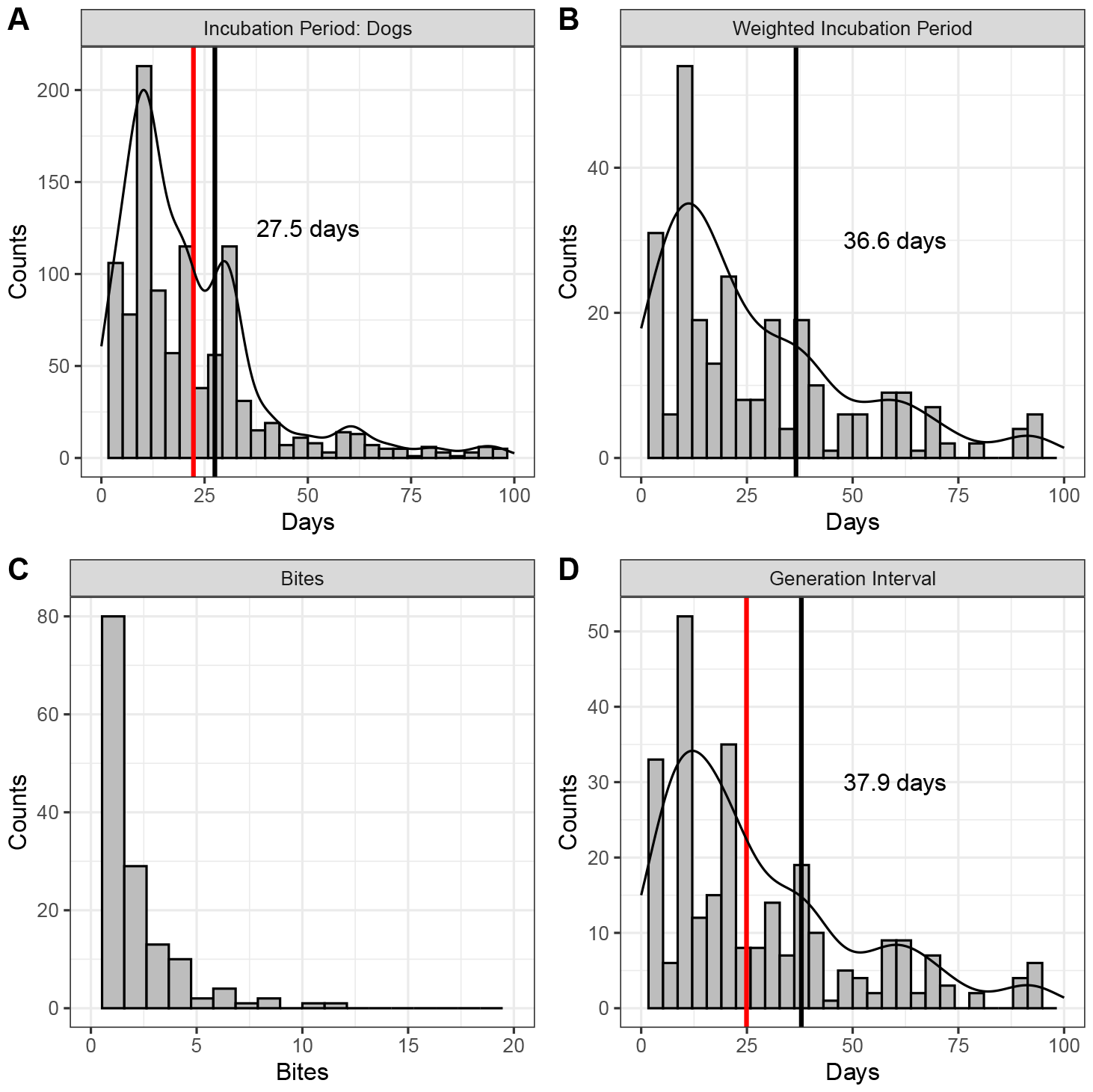
Time intervals and biting empirical distributions from contact tracing data. Panel A is the distribution of observed incubation periods. Panel B is the distribution of incubation periods weighted by each dog’s biting frequency (Panel C). The weighted distribution corresponds to the contribution of incubation periods to generation intervals (Panel D). Black vertical lines show the means of each time-interval distribution; red vertical lines show the mean incubation period and generation interval (22.3 and 24.9 days, respectively) reported by Hampson et al. (2009).

We estimated *r*_0_ from historical outbreak data (Figure 3). For a direct comparison of the method used in (Hampson et al., 2009), we also estimated *r*_0_ from an exponential model. Both methods (exponential and logistic) were applied to all phases of the global historical outbreaks. Overall, *r*_0_ estimates from the logistic model are larger with wider confidence intervals compared to *r*_0_ estimates from the exponential model (as used in (Hampson et al., 2009)).

**Figure 3:**
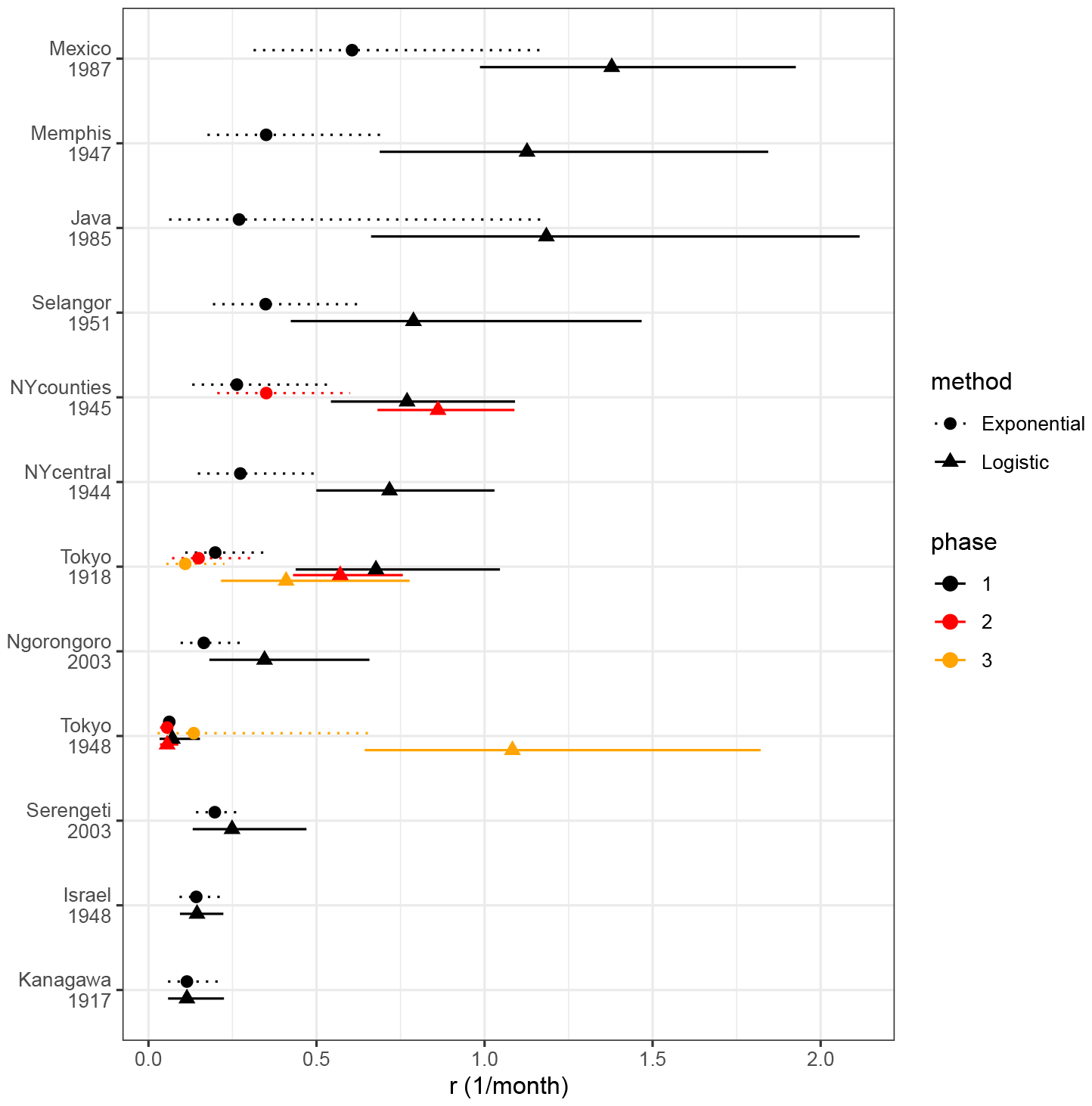
Growth rate estimates for global historical outbreaks of rabies. Estimates and 95% confidence intervals of *r*_0_ in global historical outbreaks estimated from exponential (dotted) and logistic (solid) model fits. Different colors represent different phases from the times series data.

We combined our estimates of *r*_0_ from the logistic model with the empirical *G* from our detailed Tanzanian data to produce *ℛ*_0_ estimates. Of the listed historical outbreaks, four occurred in locations with prior rabies vaccination coverage: Memphis and Shelby County, Tennessee, US (“Memphis”: 1947, 10% vaccine coverage); Serengeti, Tanzania (2003, 20% coverage); Ngorongoro, Tanzania (2003, 20% coverage); and Sultan Hamad, Kenya (1992, 24% coverage). Figure 4 shows the *ℛ*_0_ estimates using various approaches along with estimates from Hampson et al. (2009). Our estimates of *ℛ*_0_ using the logistic model and corrected *G* are larger than those previously reported (Hampson et al., 2009), with 3 locations (Java, Memphis, and Mexico) having *ℛ*_0_ greater than 2. The hybrid approach provides larger values of *ℛ*_0_ and wider confidence intervals after propagating uncertainty from both *r*_0_ and generation interval distributions with upper confidence limits greater than 2 for most locations.

**Figure 4:**
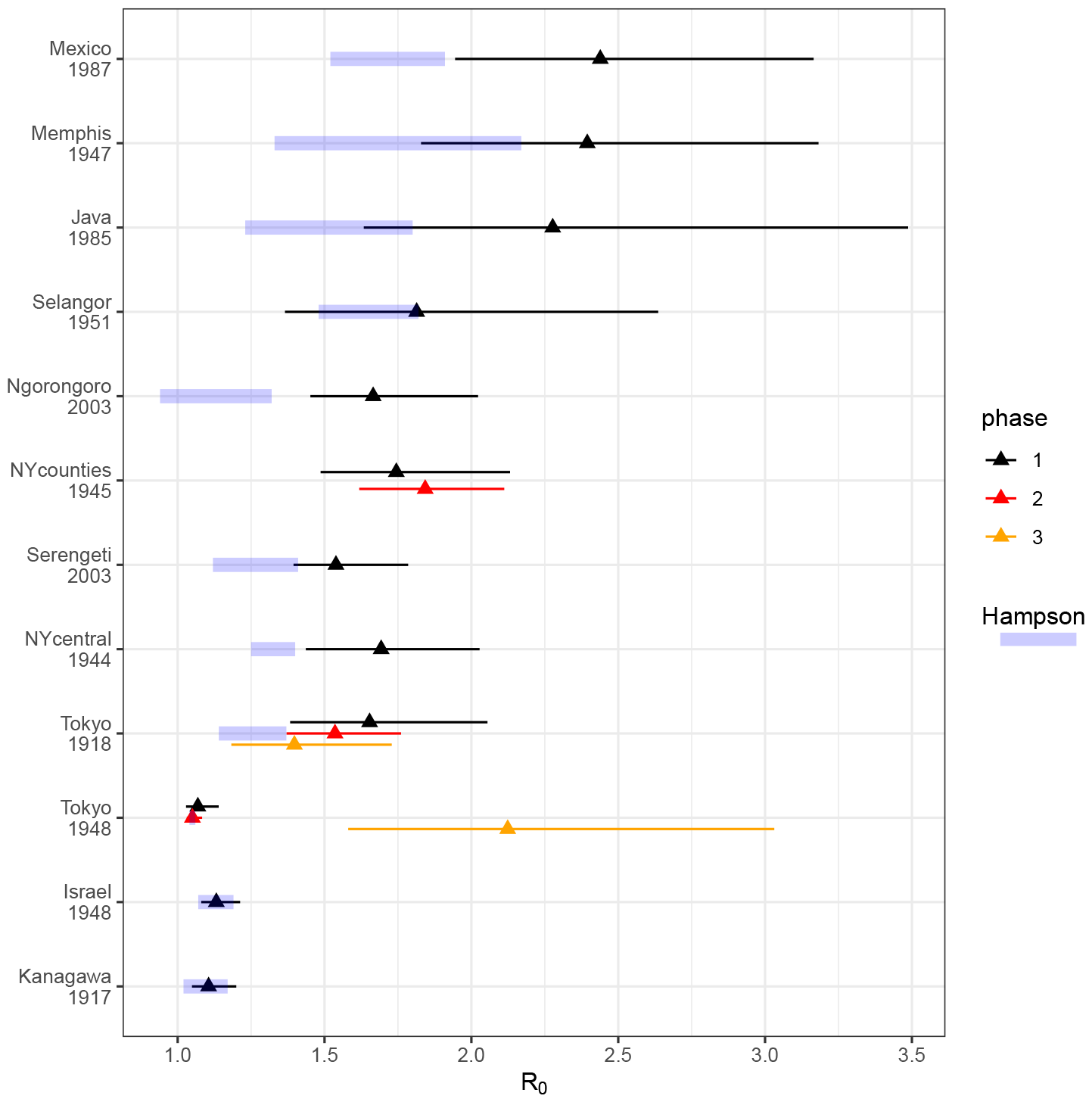
Reproductive number estimates for global historical outbreaks of rabies. Previous estimates of *ℛ*_0_ are shown in blue highlights; *ℛ*_0_ estimates and confidence interval (95% quantiles from the estimated *ℛ*_0_ sample) from our hybrid approach using the logistic model. *ℛ*_0_ values are corrected for vaccination coverage.

Lastly, we compare the effects of different estimation techniques of *r*_0_ and *G* on estimates of *ℛ*_0_ (Figure 5). For illustrative purposes, we used the 1987 outbreak in Mexico where there was no vaccination. Propagating uncertainty from both *r*_0_ and *G* generally leads to wider confidence intervals compared to previous *ℛ*_0_ estimates in Hampson et al. (2009). The *ℛ*_0_ estimate increases when we estimate *r*_0_ via the logistic model, or when we sample the full distribution of *G*, rather than plugging in the naively constructed interval as in Hampson et al. (2009). Combining the two corrections (in *r*_0_ and *G*) boosts the *ℛ*_0_ estimates even more, with even wider confidence intervals.

**Figure 5:**
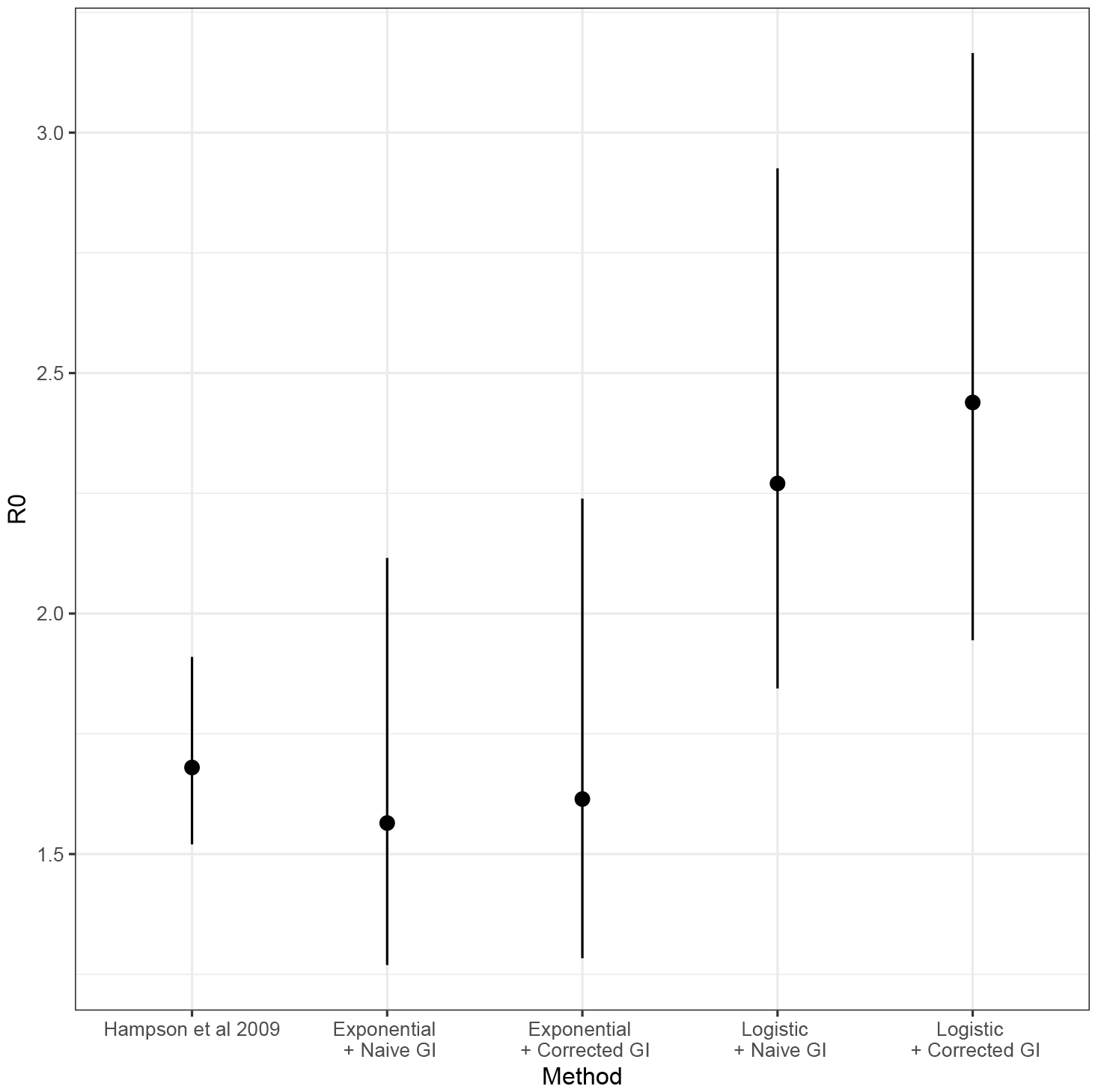
Effects of *r*_0_ estimation methods and corrected *G* on the estimates of *ℛ*_0_ in Mexico outbreak. “Exponential” represents a fitting method similar to that used by Hampson et al. (2009), but using our algorithmic windowing selection; “Logistic” uses a logistic model instead. “Naive GI” uses the *G* estimates from Hampson et al. (2009); “Corrected GI” uses the resampling method described above. Both switching from exponential to logistic fitting, and using the corrected *G* estimate, lead to increases in the estimated *ℛ*_0_. Propagating the uncertainty of *r*_0_ and *G* estimates increases uncertainty in *ℛ*_0_.

## Discussion

Our study helps to explain why rabies persists despite estimates of *ℛ*_0_ from historical outbreaks being consistently low, by showing that revised higher estimates are compatible with historical outbreak data. Here, we reanalyzed historical rabies epidemics with improved model assumptions and uncertainty propagation, showing that historical estimates of *ℛ*_0_ were downward biased and overconfident.

The basic reproductive number, *ℛ*_0_, is commonly used to summarize the risk of infectious disease and to inform control measures. Here, we used a relatively simple approach to estimating *ℛ*_0_ by combining initial growth rate (*r*_0_) estimates from incidence data and generation intervals from contact tracing data. By assuming rabies generation intervals are similar across time and space, this method allows us to combine generation intervals from the detailed Tanzania contact tracing data with growth rates estimated from incidence data from various regions across the globe. We improved on earlier work by correcting for curbed epidemic growth when estimating *r*_0_, and by developing an approach to propagate uncertainty from both *r*_0_ and *G*, resulting in higher *ℛ*_0_ estimates with wider confidence intervals.

Estimates of *ℛ*_0_ are strongly affected by estimates of the growth rate during the initial phase of the epidemic. The logistic model gives a better approximation of the initial phase of the epidemic resulting in a larger estimate of *r*_0_ compared to the exponential model (Ma et al., 2014). Our estimates of *r*_0_ account for observation error (measurements may not perfectly match reality), but not for process error (the fundamental stochasticity of the system itself). Thus we may still be underestimating the uncertainty in *r*_0_, and hence in *ℛ*_0_ (King et al., 2015). Likewise, our approach may overestimate *ℛ*_0_ since epidemic growth models can only be applied to distinct outbreaks rather than to stuttering transmission chains that are typical for diseases with low *ℛ*_0_ like rabies. Our approach also does not account for uncertainties that arise from choices about window selection, including which “phases” of outbreaks are included at all.

Re-analysis of these data also allowed us to identify an overlooked fact about rabies generation intervals: observed generation intervals are longer, on average, than intervals constructed by naively adding incubation periods and waiting times, because of within-individual correlations in time distributions and biting behaviour. The unexpected importance of these correlations could have implications for other infectious disease analyses that depend on the generation interval, as such correlations can bias the estimation of generation intervals, as shown in this study. Further investigation of how these correlations affect the overall dynamics of rabies is warranted.

In any case, our estimates suggest that the *ℛ*_0_ of rabies is larger, and more uncertain, than previously estimated. This finding may explain some of the formerly unexplained variations in the success of rabies-control programs (e.g., low levels of coverage (30–50%) have succeeded in some settings while high coverage 75% was insufficient to control rabies in others (Eng et al., 1993)). Nonetheless, our revised *ℛ*_0_ estimates still suggest that coverage required to control rabies should be feasible even in settings where *ℛ*_0_ is relatively high and that this should not be abarrier to initiating large-scale dog vaccination required for elimination.

While our primary goal was to understand why estimates of rabies *ℛ*_0_ were small with narrow confidence intervals, our analysis also revealed an interesting biological process through the lens of generation intervals from contact tracing data: the need to account for biting behaviour in the incubation period distribution, in order to match the generation interval distribution.

*ℛ*_0_ is typically used as a first approximation for interventions such as vaccination to determine herd immunity thresholds. However, both heterogeneity in contacts and the correlations between incubation periods and transmission that we observed here through the generation interval suggest that simple *ℛ*_0_ estimation methods should be used with caution. Rabies is particularly useful for exploring this effect because transmission events and latent periods are directly observable via contact tracing. The correlation effect highlighted here is likely to apply in other disease systems, but hard to detect because generation intervals are so rarely directly observable.

## Notes

### Competing Interest Statement

The authors have declared no competing interest.

## References

Abela-Ridder, B., L. Knopf, S. Martin, L. Taylor, G. Torres, and K. De Balogh (2016). 2016: the beginning of the end of rabies? The Lancet Global Health 4 (11), e780–e781.

Castillo-Neyra, R., A. M. Toledo, C. Arevalo-Nieto, H. MacDonald, M. De la Puente-León, C. Naquira-Velarde, V. A. Paz-Soldan, A. M. Buttenheim, and M. Z. Levy (2019, August). Socio-spatial heterogeneity in participation in mass dog rabies vaccination campaigns, Arequipa, Peru. PLOS Neglected Tropical Diseases 13 (8), e0007600.

Champredon, D. and J. Dushoff (2015). Intrinsic and realized generation intervals in infectious-disease transmission. Proceedings of the Royal Society B: Biological Sciences 282 (1821), 20152026.

Chowell, G. (2017). Fitting dynamic models to epidemic outbreaks with quantified uncertainty: A primer for parameter uncertainty, identifiability, and forecasts. Infectious Disease Modelling 2 (3), 379–398.

Eng, T., D. Fishbein, H. Talamante, D. Hall, G. Chavez, J. Dobbins, F. Muro, J. Bustos, M. De Los Angeles Ricardy, A. Munguia, et al. (1993). Urban epizootic of rabies in Mexico: epidemiology and impact of animal bite injuries. Bulletin of the World Health Organization 71 (5), 615.

Evans, M., J. B. Bailey, F. Lohr, W. Opira, M. Migadde, A. Gibson, I. Handel, M. Bronsvoort, R. Mellanby, L. Gamble, et al. (2019). Implementation of high coverage mass rabies vaccination in rural Uganda using predominantly static point methodology. The Veterinary Journal 249, 60–66.

Gibson, A. D., S. Mazeri, F. Lohr, D. Mayer, J. L. B. Bailey, R. M. Wallace, I. G. Handel, K. Shervell, M. Barend, R. J. Mellanby, et al. (2018). One million dog vaccinations recorded on mHealth innovation used to direct teams in numerous rabies control campaigns. PloS one 13 (7), e0200942.

Hampson, K., L. Coudeville, T. Lembo, M. Sambo, A. Kieffer, M. Attlan, J. Barrat, J. D. Blanton, D. J. Briggs, S. Cleaveland, et al. (2015). Estimating the global burden of endemic canine rabies. PLoS Neglected Tropical Diseases 9 (4), e0003709.

Hampson, K., A. Dobson, M. Kaare, J. Dushoff, M. Magoto, E. Sindoya, and S. Cleaveland (2008). Rabies exposures, post-exposure prophylaxis and deaths in a region of endemic canine rabies. PLoS Neglected Tropical Diseases 2 (11), e339.

Hampson, K., J. Dushoff, S. Cleaveland, D. T. Haydon, M. Kaare, C. Packer, and A. Dobson (2009). Transmission dynamics and prospects for the elimination of canine rabies. PLoS Biology 7 (3), e1000053.

Jagan, M., B. Bolker, J. Dushoff, D. Earn, and J. Ma (2024). epigrowthfit: Nonlinear Mixed Effects Models of Epidemic Growth. R package version 0.15.0.

King, A. A., M. Domenech de Cellés, F. M. Magpantay, and P. Rohani (2015). Avoidable errors in the modelling of outbreaks of emerging pathogens, with special reference to Ebola. Proceedings of the Royal Society B: Biological Sciences 282 (1806), 20150347.

Kitala, P., J. J. McDermott, P. Coleman, and C. Dye (2002). Comparison of vaccination strategies for the control of dog rabies in Machakos District, Kenya. Epidemiology & Infection 129 (1), 215–222.

Kurosawa, A., K. Tojinbara, H. Kadowaki, K. Hampson, A. Yamada, and K. Makita (2017). The rise and fall of rabies in Japan: A quantitative history of rabies epidemics in Osaka prefecture, 1914–1933. PLoS Neglected Tropical Diseases 11 (3), e0005435.

Kwoba, E. N., P. Kitala, L. Ochieng, E. Otiang, R. Ndung’u, G. Wambura, K. Hampson, and S. Thumbi (2019). Dog health and demographic surveillance survey in Western Kenya: Demography and management practices relevant for rabies transmission and control. AAS Open Research 2.

Ma, J., J. Dushoff, B. M. Bolker, and D. J. Earn (2014). Estimating initial epidemic growth rates. Bulletin of mathematical biology 76 (1), 245–260.

Macdonald, G. (1952). The analysis of equilibrium in malaria. Tropical Diseases Bulletin 49 (9), 813.

Mariner, J. C., J. McDermott, J. A. P. Heesterbeek, A. Catley, and P. Roeder (2005, July). A model of lineage-1 and lineage-2 rinderpest virus transmission in pastoral areas of East Africa. Preventive Veterinary Medicine 69 (3), 245–263.

Mazeri, S., A. D. Gibson, N. Meunier, M. Barend, I. G. Handel, R. J. Mellanby, and L. Gamble (2018). Barriers of attendance to dog rabies static point vaccination clinics in Blantyre, Malawi. PLoS Neglected Tropical Diseases 12 (1), e0006159.

Minghui, R., M. Stone, M. H. Semedo, and L. Nel (2018). New global strategic plan to eliminate dog-mediated rabies by 2030. The Lancet Global Health 6 (8), e828–e829.

Mtema, Z., J. Changalucha, S. Cleaveland, M. Elias, H. M. Ferguson, J. E. Halliday, D. T. Haydon, G. Jaswant, R. Kazwala, G. F. Killeen, et al. (2016). Mobile phones as surveillance tools: implementing and evaluating a large-scale intersectoral surveillance system for rabies in Tanzania. PLoS Medicine 13 (4), e1002002.

Nadal, D., B. Abela-Ridder, S. Beeching, S. Cleaveland, K. Cronin, R. Steenson, and K. Hampson (2022). The impact of the first year of the covid-19 pandemic on canine rabies control efforts: a mixed-methods study of observations about the present and lessons for the future. Frontiers in Tropical Diseases 3, 866811.

Park, S. W., D. Champredon, J. Weitz, and J. Dushoff (2018). Exploring how generation intervals link strength and speed of epidemics. bioRxiv, 312397.

Rupprecht, C., J. Barrett, D. Briggs, F. Cliquet, A. Fooks, B. Lumlertdacha, F. Meslin, T. Müler, L. Nel, C. Schneider, et al. (2008). Can rabies be eradicated? Developments in biologicals 131, 95–121.

Taylor, L. H., K. Hampson, A. Fahrion, B. Abela-Ridder, and L. H. Nel (2017). Difficulties in estimating the human burden of canine rabies. Acta Tropica 165, 133–140.

Tepsumethanon, V., H. Wilde, and F. X. Meslin (2005). Six criteria for rabies diagnosis in living dogs. J Med Assoc Thai 88 (3), 419–22.

Wallace, R. M., H. Reses, R. Franka, P. Dilius, N. Fenelon, L. Orciari, M. Etheart, A. Destine, K. Crowdis, J. D. Blanton, et al. (2015). Establishment of a canine rabies burden in Haiti through the implementation of a novel surveillance program. PLoS Neglected Tropical Diseases 9 (11), e0004245.

Wallinga, J. and M. Lipsitch (2006). How generation intervals shape the relationship between growth rates and reproductive numbers. Proceedings of the Royal Society B: Biological Sciences 274 (1609), 599–604.

